# RAMA: a global genome size database of invasive tetrapod species

**DOI:** 10.64898/2026.11.12.766098

**Authors:** Anna Miall, Thomas W. Bodey, Ross N. Cuthbert, Daniel Pincheria-Donoso

## Abstract

**Motivation:** Rising numbers of biological invasions have triggered widespread erosion of environmental and socio-economic stability worldwide. To predict and understand the impacts of current and future invasive non-native species (INNS), invasion science has increased collective efforts to elucidate the role of functional traits as drivers of INNS establishments and impacts.

Tetrapod vertebrates (reptiles, amphibians, mammals and birds) represent exceptionally well-studied animal lineages with broad data availability, while serving as regular model systems on the predictors of invasion success and impacts. However, while efforts exist to compile data on phenotypic traits, the availability of databases linking phenotypes and genomes remains neglected among animals. Here, we introduce the Reptilia, Amphibia, Mammalia and Aves (RAMA) database, an open-access, global-scale resource compiling genome size records for 3,539 non-native species together with invasion success and impact data. RAMA reveals relatively low coverage of tetrapod genome sizes (9.51%), as well as highlighting an imbalance in genomic and invasion data availability among taxa. Genome data is more commonly available for mammals (18.56%) and birds (16.42%) than for amphibians (5.1%) and reptiles (2.42%). RAMA, the first database to assemble animal genome size and invasion data together, encourages further investigations and incorporations of a genomic perspective within invasion science to develop predictive research and inform proactive management strategies.

***Main types of variables contained:*** species-level genome and invasion records.

***location and grain:*** Global, tetrapod-wide.

***Time period and grain:*** Extant (1883 to 2024).

***Major taxa and level of measurement:*** 3,539 tetrapod species.

***Software format:*** .csv

## 1. Background and summary

The expansion of globalisation networks has significantly enhanced the emergence of pathways for (deliberate and unintentional) the human-mediated movement of species from their native to non-native regions, leading to a sustained growth in the number of invasive non-native species (INNS)(Seebens et al., 2025). A critical consequence of these biological invasions is the aggregate detrimental impacts they exert to ecosystem dynamics, biodiversity persistence, and socio-economic welfare (Seebens et al., 2017, Diagne et al., 2021a, Carneiro et al., 2025, Carneiro et al., 2026, Courchamp et al., 2026). Given the rapidly expanding pressures arising from INNS, research efforts have been directed to the identification of the factors and mechanisms behind the process from introduction to successful invasions (Crowl et al., 2008, Hulme, 2009, Blackburn et al., 2011, Richardson and Pyšek, 2011, Diagne et al., 2021b, Haubrock et al., 2025). Alongside these efforts, global projections of future invasions have catalysed ambitious international targets to understand, predict and mitigate INNS (Seebens et al., 2021). Research has enabled the development of proactive management strategies targeting earlier stages of biological invasions to reduce overall spread and impact (Reaser et al., 2020, IUCN, 2024). Further development and improved efficiency in identifying future and high impact invaders depends on predictive research advancements and, subsequently, practical applications.

Traditionally, research efforts have focused on a wide range of functional (demographic and fitness relevant) traits to establish the role that pre-existing adaptations (i.e., traits that fit the demands of a non-native range) or potential for rapid adjustments via plasticity or adaptation (i.e., ecological generalism, fast life history pace) play in the establishment of non-native species into invasives (Capellini et al., 2015, Pepe, 2022, Cuthbert et al., 2025). In particular, positive relationships between invasion success and high fecundity traits have been observed for invasive plant species (e.g., greater seed mass, production and dispersal) and animals (e.g., increased litter/brood size and frequency) (Moodley et al., 2013, Capellini et al., 2015, Moravcová et al., 2015). Additionally, plasticity of life history traits has been shown to increase the likelihood of invasion success, through enhancing a species’ capacity to adjust to novel conditions without having to undergo an adaptive response (Richards et al., 2006, Davidson et al., 2011, Le Roux, 2021). Although, depending on the taxon and pathway for introduction, the relevance of these traits likely differs among invasion stages.

Despite these extensive efforts devoted to the elucidation of phenotypic drivers underlying invasion success, the use of genomic factors has lagged considerably behind (Pyšek et al., 2018, Guo et al., 2024, Kołodziejczyk et al., 2025). In recent years, there has been interest in addressing the role that variation in genome size plays in the likelihood of population adapting to the demands of their environments (Pincheira-Donoso et al., 2023). These studies have investigated intrinsically demographic predictions, such as the relationship between genome size and functional traits (Gregory, 2005, Lertzman-Lepofsky et al., 2019, Decena-Segarra et al., 2020, Beaudreau et al., 2021), and the influence of variation of genome size on likelihood of population extinctions (Kraaijeveld, 2010, Lertzman-Lepofsky et al., 2019, Pincheira-Donoso et al., 2023). For example, the ‘large genome constraint hypothesis’ predicts that species with large genomes are more likely to experience adverse costs such as reduced phenotypic plasticity, slower growth rates, and higher extinction risk (Pagel and Johnstone, 1992, Vinogradov, 2003, Gregory, 2005, Knight et al., 2005, Kraaijeveld, 2010, Arnqvist et al., 2015, Issei et al., 2017, Faizullah et al., 2021, Soto Gomez et al., 2024). Invasive plant populations are observed to have a tendency for small genomes, and the functional traits (greater environmental tolerance and growth rates) associated with small-genome sizes aid invasion success (Lavergne et al., 2010, Pyšek et al., 2018, Pyšek et al., 2023, Meyerson et al., 2024). Plasticity, a trait well recognised within successful invasions, is further negatively influenced by plant genome size (Richards et al., 2006, Meyerson et al., 2020). Given that the vast variation in genome size across animals has major functional and evolutionary implications, it is surprising that its role as a factor underlying invasion success has been limited to plant studies, with animal invasions remaining fundamentally neglected (Gregory, 2011, Pincheira-Donoso et al., 2023).

Assembling invasion and genome data together and expanding on genomic analysis into determining animal invasion likelihood would advance predictive invasion science and aid in management prioritisation. Here, the Reptilia, Amphibia, Mammalia and Aves (RAMA) database is presented as a global-scale resource containing genome and invasion data across four tetrapod lineages. This paper provides details on the database development, a comprehensive description of the data and practical details for future users.

## 2. Methodology

### 2.1 Data sources and collation

The Reptilia, Amphibia, Mammalia and Aves (RAMA) database is constructed from multiple, open-access data sources providing species-level invasion and genome size data in a single comprehensive database, and a standardised taxonomic classification of the species (Fig. 1, Table 1). The initial step of database construction was to obtain a (native/non-native) species checklist for each taxonomic class, followed by genome and invasion data extraction (Fig. 1, Table 1).

**Figure 1.**
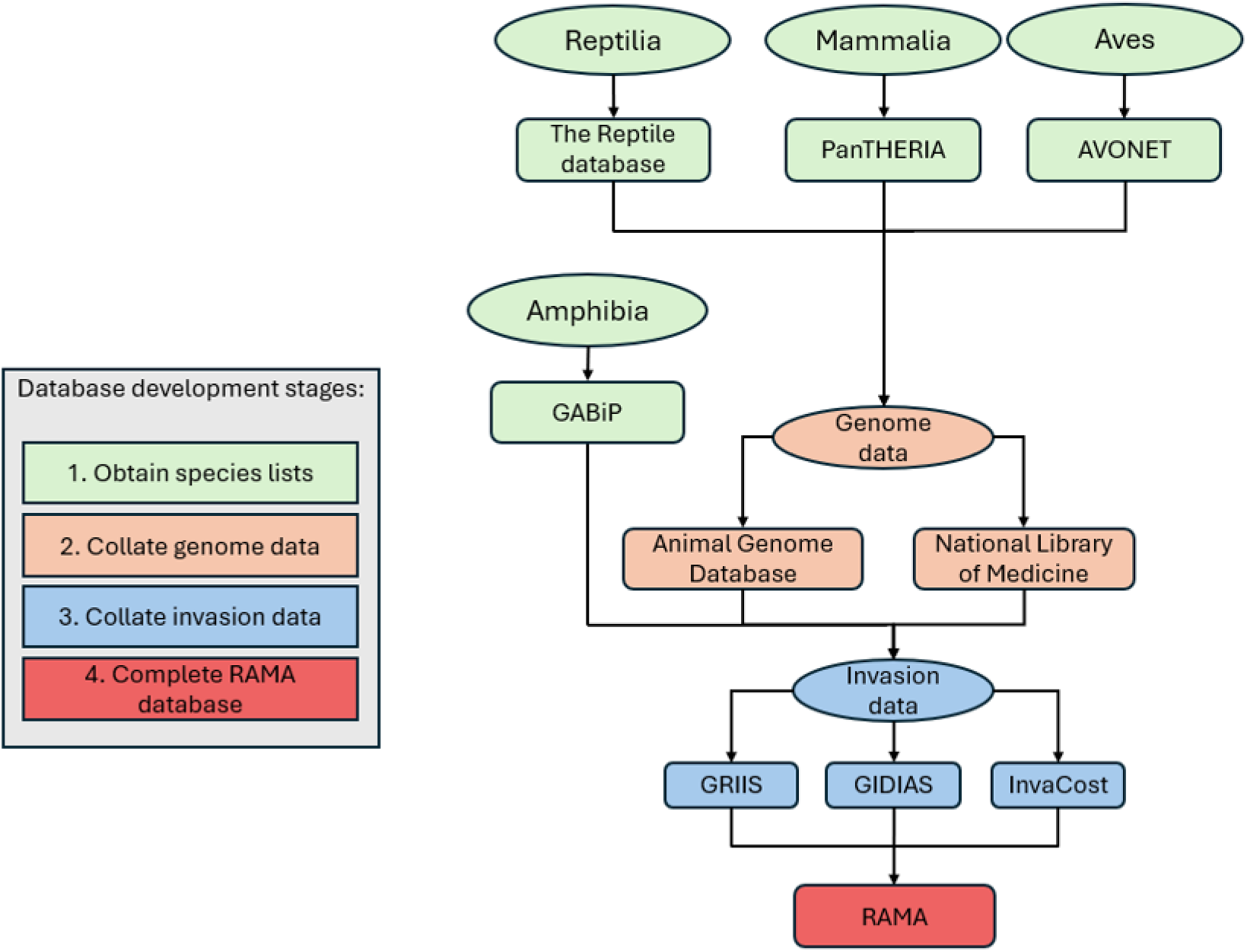
The development stages used to construct the Reptilia, Amphibia, Mammalia and Aves (RAMA) database, illustrating the structured process of data collection and the data source utilised.

**Table 1.** Structure of the sources of data collected and collated for the Reptilia, Amphibia, Mammalia and Aves (RAMA) database, with details specifying the format (species lists, genome and invasion data), the source, as well as the type of data provided by each database. Further detail is provided in Supplementary Table 2.

| Data type | Database | Data provided | Variable description | Variables format | Access date (mm/yy) |
|---|---|---|---|---|---|
| **Species list** | The Reptile Database (Uetz et al., 2026) | Reptilia class species list | The taxonomic classification of the reptile species, from class- to species-level | Character | 05/25 |
|  | Global Amphibian Biodiversity Project (GABiP) | Amphibia class species list | The taxonomic classification of the amphibian species, from class- to species-level | Character | 06/25 |
|  | PanTHERIA (Jones et al., 2009) | Mammalia class species list | The taxonomic classification of mammal species, from class- to species-level | Character | 04/25 |
|  | AVONET (Tobias et al., 2022) | Aves class species list | The taxonomic classification of the avian species, from class- to species-level | Character | 01/25 |
| **Genome** | GABiP | Amphibian genome size (c-value) | The median genome size value of an amphibian species | Numeric continuous | 06/25 |
|  | The Animal Genome Database (Gregory, 2025) | Genome size (c-value) | The median genome size value of species for reptiles, birds and mammals | Numeric continuous | 01/25 to 06/25 |
|  | National Library of Medicine (NCBI, 2025) | Genome size (pb and mb) | The median genome size value of species for reptiles, birds and mammals | Numeric continuous | 01/25 to 06/25 |
| **Invasion** | Global Impacts Database of Invasive Alien Species (GIDIAS) (Bacher et al., 2025) | Invaded locations | Records of the locations (geographic regions and land types) an INNS are present, structured as occurrence by each regional/land type category (e.g., invaded_Africa) | Binary | 07/25 |
|  |  | Invaded habitats | The habitat type the INNS are present within, structured by occurrence in each habitat type (e.g., invaded.Marine) | Binary |  |
|  |  | Invasion spatial scale | Records of the spatial scale of the invasions of the INNS, structured by either local or global spatial scales (e.g., local.spatial.scale) | Binary |  |
|  |  | Impact occurrence | The type of impact INNS’s (nature or nature’s contribution to people) including direction of impact (positive, negative or neutral) (e.g. Nature.negative.impact) | Binary |  |
|  |  | Mechanisms of impact | The mechanism in which an INNS nature impacts are inflicted, structured by use of each mechanism type (e.g., competition.Nature) | Binary |  |
|  |  | Mean magnitude of impact | The average ‘magnitude of impact’ score associated with an INNS | Ordinal |  |
|  |  | First and last year of impact | The first and last year an INNS impacts were recorded | Numeric discrete |  |
|  | InvaCost (Diagne, 2020) | Locations of impact | The land types (urban, protected area and island) or geographic regions in which an INNS economic impacts are recorded, structured as occurrence by each regional/land type category (e.g., InvaCost_PA_invaded) | Binary | 03/25 |
|  |  | Probable starting & ending years of economic impact | The estimated starting and ending years of the recorded economic impacts | Numeric continuous |  |
|  |  | Occurrence type | Records indicating whether the economic impacts are on-going or were a one-time occurrence, structured by each occurrence type (e.g., InvaCost_Occurrence_Onetime) | Binary |  |
|  |  | Mean cost of INNS | The average monetary cost of the impacts caused by INNS | Numeric continuous |  |
|  |  | Total cost of INNS | The total monetary cost of the impacts caused by INNS | Numeric continuous |  |
|  |  | Sectors impacted | Records of which sectors were economically impacted by the INNS, structured by each sector impacted (e.g., inva_impacted_Agriculture) | Binary |  |
|  |  | Cost type | The type of costs (damage and management) generated by the INNS, structured by each cost type (e.g., inva_CostType_Damage) | Binary |  |
|  |  | Management type | The type of management (knowledge, pre-invasion and post-invasion) used against the INNS, structured by each management type (e.g., inva_ManagmentType_Knowledge) | Binary |  |
|  | Global Register of Introduced and Invasive Species (GRIIS) (Pagad et al., 2018, Pagad et al., 2022) | Establishment status | A record as to whether the species is recognised as an established non-native species (Establishment_Status) | Binary | 08/25 |
|  |  | Species invasive status | A record as to whether the species is recognised as an invasive species | Categorical |  |
|  |  | Number of countries invaded | The total number of countries where a species is known to be established. | Numeric count |  |
|  |  | Number of continents invaded | The total number of continents where a species is known to be established. | Numeric count |  |

### 2.2 Genome and invasion records

The Global Amphibian Biodiversity Project (GABiP) database compiles morphological, life history and genome size data for the world’s amphibian species (Pincheira-Donoso et al., 2021, Johnson et al., 2023, Pincheira-Donoso et al., 2023, de Oliveira Caetano et al., 2025). For the remaining classes, genome records were extracted from the ‘Animal Genome Database’ (AGD) and ‘National Centre for Biotechnology Information’ (NCBI) genome database (Gregory, 2025, NCBI, 2025). The AGD, consisting of records sourced only from published data, was selected as the primary database due to high reliability, with NCBI used when data were only present in this source (more detail provided within Supplementary Material).

The invasion data include invasion success (establishment records, numbers of countries and continents established within) as well as the associated occurrence and magnitude of negative impacts caused by said species (ecological and economic impact records) (Table 1). Invasion data were sourced from the Global Impacts Database of Invasive Alien Species (GIDIAS (Bacher et al., 2025)), InvaCost (Diagne et al., 2020) and the Global Register of Introduced and Invasive Species (GRIIS, (Pagad, 2022))(Table 1, see Supplementary Material). GIDIAS provided invasion data regarding invaded location, invaded habitat, and invasion spatial scale. Invasion data were expanded from categorical, invasion event-level records into binary, species-level variables to retain all information from the records while developing a simple and tractable dataset for analysis. For example, InvaCost impacted sector data (Agriculture, Authorities-Stakeholders, Environment, Fishery, Forestry, Health, Public and social welfare and Unspecified) were converted from invasion-level, categorical format into a binary, species-level format (example; inva_impacted_Agriculture, inva_impacted _ AuthoritiesStakeholders, etc). For a more detailed description of the RAMA construction and variables, see the Supplementary Methods in Supplementary Material.

### 2.3 Analyses

Data analyses were performed in R studio version 2025.09.1 (R_Core_Team, 2025). Packages ‘*ggplot2’* and ‘*dyplr’* were used to prepare the database and records for analysis (Wickham, 2016, Wickham et al., 2023). Two chi-squared tests of independence were performed to assess differences in the availability of genome and establishment data as categorical responses (Has Value or No Value), with taxonomic class as the predictors (R_Core_Team, 2025). To determine whether there were statistically significant differences between 1) genome size, 2) number of countries invaded, and 3) impact magnitude across tetrapod classes, Kruskal-Wallis tests (R package ‘stats’) were performed and followed by Dunn’s post-hoc tests where necessary (R package ‘FSA’, (Ogle and Ogle, 2017, R_Core_Team, 2025). Accounting for smaller sample sizes, a two-sided Fisher’s Exact test was performed to assess the differences in recorded impact types (e.g., Ecological, Economic, Both and None) as a categorical response and the four taxonomic classes as the predictor, followed by a pairwise Fisher’s exact test of independence. All post hoc tests used Bonferroni corrections. Effect sizes were calculated using r package ‘rstatix’ (Kassambara, 2019).

## 3. Results and Discussion

### 3.1 Data usage

The RAMA database is provided in a single, long-format spreadsheet containing genome and invasion data records across four taxonomic classes for 3,539 species (‘RAMA_df.csv’). A variable description and the data sources are provided for all records in an additional spreadsheet (‘RAMA_metadata.csv’) (see Supplementary Material).

### 3.2 Genome data availability and range

In RAMA, percentage coverage of genome size records (relative to total species diversity within each class) differs among mammals (18.56%), birds (16.42%), amphibians (5.1%) and reptiles (2.42%), with an overall left-skew in genome size across taxa but with large-genome amphibian outliers (Fig. 2). Genome size data availability varied significantly among all classes (Table 2, see Supplementary Material). The RAMA data coverage is taxonomically asymmetric, with much higher representation of mammals and birds, relative to the lower representation of amphibian and reptile species (Hughes et al., 2021, Seebens et al., 2025). Variation of genome size across RAMA extends from 0.08 to 120.57 c-value (Table 3, Fig. 2), and genome size was found to be significantly different among classes (Table 2); furthermore, the Dunn’s post hoc indicated that significant differences were present between each taxonomic group (Table 2). Despite the low coverage, the amphibian and reptile species within RAMA cover the documented overall genome size range for both lineages from 0.95 to ∼120 pg and 1.09 to 5.44 pg respectively (Table 3) (Gregory, 2011).

**Figure 2.**
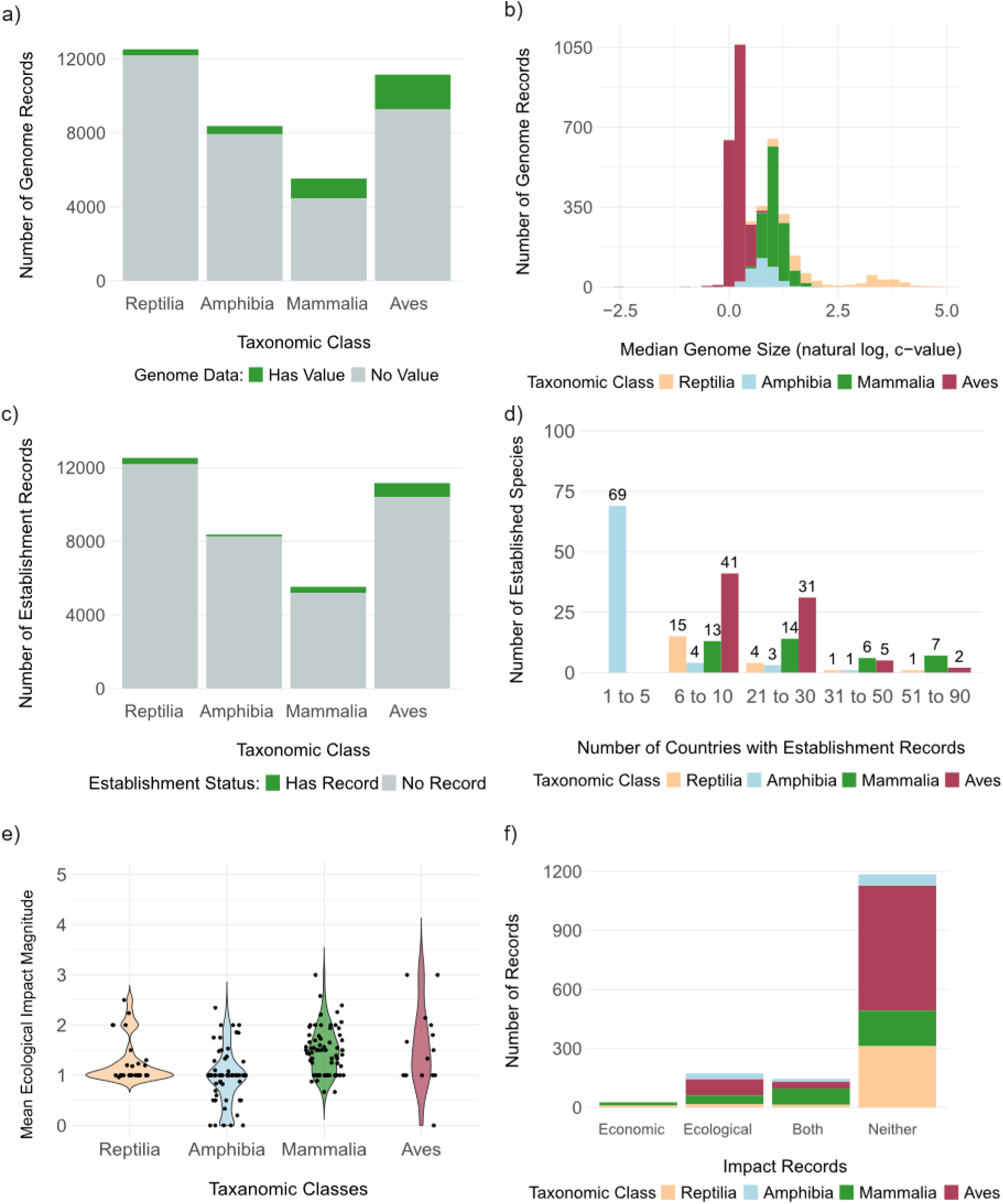
The distribution of: **a)** coverage of recorded genome sizes for species; reptiles with 2.42%, amphibians with 5.1%, mammals with 18.56% and birds with 16.42%, **b)** median genome sizes (c-value) in a natural logarithm, **c)** establishment records of 1212 species (3.26%) species; establishment records are provided for 271 reptile species (2.18%), 77 amphibians (0.92%), 223 mammal species (4.11%) and 641 avian species (5.83%), **d)** establishment records across binned numbers of countries, **e)** mean ecological impact magnitudes, and **f)** the impact record types (economic, ecological, both or neither) separated by taxonomic classes Reptilia, Amphibia, Mammalia and Aves.

**Table 2.** Outputs from the statical analysis of RAMA records, including X^2^ and p-values of Pearson’s chi-squared, Kruskal-Wallis and Fisher’s exact tests with associated effect sizes and pairwise comparisons.

| **Statical test** | **Response** | **Outputs** |  | **Effect size** | **Pairwise comparison** | **p-value** |
|---|---|---|---|---|---|---|
|  |  | X<sup>2</sup> | p-value |  |  |  |
| **Pearsons chi-squared** | Genome data availability | 2041.4 | **<0.001** | 0.17 | **Amphibia – Aves** | **< 0.001** |
|  |  |  |  |  | **Amphibia - Mammalia** | **< 0.001** |
|  |  |  |  |  | **Aves - Mammalia** | **< 0.001** |
|  |  |  |  |  | **Amphibia - Reptilia** | **< 0.001** |
|  |  |  |  |  | **Aves - Reptilia** | **< 0.001** |
|  |  |  |  |  | **Mammalia - Reptilia** | **< 0.001** |
|  | Establishment data availability | 432.78 | **<0.001** | 0.17 | **Amphibia – Aves** | **< 0.001** |
|  |  |  |  |  | **Amphibia - Mammalia** | **< 0.001** |
|  |  |  |  |  | **Aves - Mammalia** | **< 0.001** |
|  |  |  |  |  | **Amphibia - Reptilia** | **< 0.001** |
|  |  |  |  |  | **Aves - Reptilia** | **< 0.001** |
|  |  |  |  |  | **Mammalia - Reptilia** | **< 0.001** |
| **Kruskal-Wallis** | Genome size variation | 2758.5 | **<0.001** | 0.07 | **Amphibia – Aves** | **< 0.01** |
|  |  |  |  |  | **Amphibia - Mammalia** | **< 0.001** |
|  |  |  |  |  | **Aves - Mammalia** | **< 0.01** |
|  |  |  |  |  | **Amphibia - Reptilia** | **< 0.001** |
|  |  |  |  |  | **Aves - Reptilia** | **< 0.001** |
|  |  |  |  |  | **Mammalia - Reptilia** | **< 0.001** |
|  | Number of countries invaded | 20.5 | **<0.001** | <0.01 | Amphibia – Aves | 0.96 |
|  |  |  |  |  | Amphibia - Mammalia | 0.42 |
|  |  |  |  |  | Aves - Mammalia | 1.00 |
|  |  |  |  |  | Amphibia - Reptilia | 1.00 |
|  |  |  |  |  | **Aves - Reptilia** | **< 0.001** |
|  |  |  |  |  | **Mammalia - Reptilia** | **< 0.001** |
|  | Mean impact magnitude | 33.9 | **<0.001** | 0.03 | Amphibia – Aves | 0.12 |
|  |  |  |  |  | Amphibia - Mammalia | 0.31 |
|  |  |  |  |  | **Aves - Mammalia** | **<0.001** |
|  |  |  |  |  | Amphibia - Reptilia | 1.00 |
|  |  |  |  |  | **Aves - Reptilia** | **< 0.01** |
|  |  |  |  |  | Mammalia - Reptilia | 1.00 |
| **Fisher’s Exact** | Impact type | NA | **<0.001** | 0.21 | **Amphibia – Aves** | **< 0.001** |
|  |  |  |  |  | **Amphibia - Mammalia** | **< 0.001** |
|  |  |  |  |  | **Aves - Mammalia** | **< 0.001** |
|  |  |  |  |  | **Amphibia - Reptilia** | **< 0.001** |
|  |  |  |  |  | **Aves - Reptilia** | **< 0.001** |
|  |  |  |  |  | **Mammalia - Reptilia** | **< 0.001** |

**Table 3.** The Reptilia, Amphibia, Mammalia and Aves (RAMA) database genome size (pg) variation displayed across quantiles (0%, 25%, 50%, 75% and 100%) and the mean for all species within RAMA as well as for each taxonomic class.

| **Taxon** | **Quantiles (%)** |  |  |  |  | **Mean** |
|---|---|---|---|---|---|---|
|  | 0 | 25 | 50 | 75 | 100 |  |
| **RAMA** | 0.08 | 1.23 | 1.58 | 2.78 | 120.57 | 3.82 |
| **Reptilia** | 1.09 | 1.81 | 2.11 | 2.58 | 5.44 | 2.23 |
| **Amphibia** | 0.95 | 3.9 | 6.18 | 28.16 | 120.57 | 17.82 |
| **Mammalia** | 1.63 | 2.45 | 2.76 | 3.27 | 8.4 | 2.95 |
| **Aves** | 0.08 | 1.12 | 1.23 | 1.33 | 2.35 | 1.25 |

### 3.3 Establishment and impact biases

From the 1212 species within RAMA with genome size and a positive establishment record, RAMA revealed low percentage coverage of establishment data across all four lineages (amphibians: n = 77, 0.92 %, mammals: n = 223, 4.11%, reptiles: n = 272, 2.18%, birds: n = 641, 5.83%) (Fig. 2); significant differences in coverage were found across all pairwise comparisons (Table 2). Regarding the number of countries established within, our Kruskal-Wallis provided evidence of significant difference between taxonomic classes; a Dunn’s test indicated that significant differences were present between Aves and Reptilia as well as Mammalia and Reptilia (Table 2). Mean ecological impact magnitude significantly differed among classes (Table 2), with post-hoc tests revealing significant differences present between bird and mammal and bird and reptile species (Table 2). Impact type was found to vary significantly across tetrapod classes, these differences were proposed to be present among all pairwise comparisons (Table 2). Our findings provide further support for asymmetry in invasion impact data regarding taxa, region and habitat type (McGeoch et al., 2010, Thomsen, 2014, Cuthbert et al., 2021).

These findings could represent the taxonomic influence of invasion pathways, since the principal routes of introduction differ among the tetrapod classes. The international pet trade is a prominent source of INNS through accidental escapees and purposeful releases for avian, reptilian and amphibian species (Kraus, 2009, Carpio et al., 2020). In contrast, purposeful introductions for biocontrol, hunting and agriculture purposes are the most common routes for mammalian invaders (Blackburn et al., 2017, Carpio et al., 2020). The number of countries invaded is also strongly influenced by variation in invasion pathways, propagule and colonisation pressures globally (Soto et al., 2026), with the number and frequency of introductions differing substantially among introduction pathways and impacting on establishment success (Lockwood et al., 2005). While our assessment of number of countries invaded does not consider geographic spread of the invasions within nations, our findings coarsely suggest invasion spread varies by RAMA classes. Indeed, previously observed vertebrate establishment and spread variations are thought to stem from differing drivers of success across species lineages (Allen et al., 2013).

## 4. Conclusion

We introduce the RAMA database as the first tetrapod, species-level data source for combined invasion records and genome sizes, with 3,539 species, with accompanying analyses revealing low genome data coverage and strong geographic and taxonomic biases across invasion records. The analysis reveals persisting research disparity and potential for biases to be distorting the assessment of INNS abundance, spread and level of threat via over and under representation. Moreover, RAMA provides a unique data source where species genome sizes have been aligned with invasion records to expand future predictive trait research. Future RAMA iterations could expand across broader and unexplored lineages as well as including additional genomic metrics.

## Supporting information

RAMA_metadata.csv

RAMA_df.csv

