## Supplementary Material for "RAMA: a global genome size database of invasive tetrapod species"

### 1. Supplementary Methods

#### 1.1 Genome records

Species names (*Genus species*) from species lists were manually inputted into the AGD. If a single genome value was provided for a species, the value was recorded as minimum, maximum and median genome size. When multiple values were provided, the lowest value was recorded as the minimum genome size, the highest value was recorded as the maximum value and the median was calculated from the minimum and maximum values. When no records were recovered from the AGD, the search was repeated within the NCBI database following the same process. The NCBI database recorded whether a record was from a 'reference' or 'contaminated' genome sequence: 'referenced' records refer to records which are single representatives for said species; and 'contaminated' status indicated the genome sequence was exposed to and potentially contain content from other organisms. Considering potential misclassification of contaminated genomes, no 'contaminated' records were used (Lupo et al., 2021). When the NCBI database provided multiple records, only 'referenced' values were extracted for data quality assurance. However, when 'referenced' values were unavailable and non-referenced values were provided, the median, minimum and maximum genomes size were calculated from the samples provided. NCBI genome sizes are expressed in basepairs (Gb) so were converted into c-values (picograms) to align with AGD data (Doležel et al., 2003).

Species check lists and genome data extraction were conducted manually on Excel. All further database developments and data analysis were conducted in R (R 3.6.0) through R studio version 2025.05.0+496 (Team, 2025). When combining the data from disparate sources, GBIF and the linked R package '*taxize*' were used in tandem to resolve any misclassification (Chamberlain et al., 2020).

#### 1.2 Invasion records

Establishment success is defined here as a species recorded as self-sustaining, non-native population. Establishment success records were sourced from the GRIIS database (Pagad et al., 2018, Pagad et al., 2022). Within GRIIS, species establishment data fall under 'establishmentMeans' with three categories; 'Alien' (a species moved beyond the native range), 'Native-Alien' (a species native to area of a country but introduced into an area outside of its native range) and 'Cryptogenic/Uncertain' (species origin unknown) (see Table S1, (Pagad et al., 2018)). Species were only considered as successfully established when categorised as 'Alien' or 'Native-Alien'.

Invasion impacts were divided into ecological and economic. Ecological impacts were defined in line with GRIIS, as when a species introduction or spread were recognised as threats to biological diversity (Table S2). Ecological impact status was collected from the GRIIS database where species were recorded as 'Invasive' under 'isInvasive'; this decision was made since the GRIIS criteria for an invasive species is for a species introduction to threaten biological diversity. Economic impact records were collected from InvaCost (Table S2). Impact records were integrated into RAMA as binary values, where species with recorded ecological and/or economic impact records are recorded as '1' (Table S2). An additional column 'Impact Type' clarifies whether species have 'No Impact', 'Ecological', 'Economic' or 'Both'. The economic impact monetary values (total and mean) were further calculated from all recorded values for a species from InvaCost data under 'Raw_cost_estimate_2017_USD_exchange_rate' using only 'observed' entries of 'high' reliability (Table S2). The InvaCost database provided records which cover the economically impactful invasion location (geographic region, habitat type), earliest starting and latest ending year, occurrence type (ongoing or onetime), impacted sector, cost type and management type (Diagne et al., 2020b)(see Supplementary Materia). These InvaCost records were expanded into binary, dummy variables within RAMA (Table S2).

GIDIAS provided records on invasion impacts; including geographic regions, habitat types, the spatial scale, type of impact, impact magnitude as well as records of the first and last impact years (Bacher et al., 2025). The categorical GIDIAS data are recorded in RAMA as binary dummy variables. For each invasion record, the GIDIAS database provided an EICAT impact magnitudes for each invasion record which is scaled from 1 to 5 (Bacher et al., 2025). Due to the invasion- to species-level conversion between databases, a mean value was calculated from each recorded EICAT impact magnitudes for a single species and one mean impact magnitude value was assigned to a species. Regarding the ecological and economic impact year records, the earliest first impact and the latest last impact year were assigned to each species.

***Table S1 -*** *Terms and definitions taken from and used within the Global Register of Introduced and Invasive Species and used throughout the development of the Reptilia, Amphibia, Mammalia and Aves database (Pagad et al., 2018).*

| Term | Definition |
|---|---|
| Alien/Introduced | A species, subspecies or (for plants) variety or cultivar, moved by human activities beyond the limits of its native geographic range, or resulting from breeding or hybridization and being released into an area in which it does not naturally occur, and includes any part, gametes or propagule of such species that might survive and subsequently reproduce. |
| Cryptogenic | Species of unknown biogeographic history that cannot be ascribed to being native or alien. |
| Uncertain | Species recognised as clearly alien although their specific geographic origin is unknown. |
| Invasive | A taxon whose introduction and/or spread threatens biological diversity (Convention on Biological Diversity; http://www.cbd.int/decision/cop/). |
| Native-Alien | Species native to some areas of a country or territory but introduced by humans into places outside of their natural range of distribution in that country, where they become established and disperse. |

### InvaCost format

| Genus_species | Impacted_sector | Type_of_cost |
|---|---|---|
| Vulpes vulpes | Agricultural | Damage |
| Vulpes vulpes | Authorities-Stakeholders | Management |
| Trachemys scripta | Authorities-Stakeholders | Management |
| Trachemys scripta | Authorities-Stakeholders | Management |

### RAMA format

| Genus_species | Impacted_Agriculture | Impacted_Authorities-Stakeholders | Damage_Cost | Management_cost |
|---|---|---|---|---|
| Vulpes vulpes | 1 | 1 | 1 | 1 |
| Trachemys scripta | 0 | 1 | 0 | 1 |

***Figure S1*** *- Visual example of workflow used to convert data into binary values within the Reptilia, Amphibia, Mammalia and Aves database*

***Table S2 –*** *The variables of the Reptilia, Amphibia, Mammalia and Aves (RAMA) database, with the variable names as appearing in the RAMA database, definitions, metrics and original data sources.*

| Column Name | Description | Value | Sourced by |
|---|---|---|---|
| Species | The taxonomic classification of the species, at a species-level and formatted as 'Genus species' | Character | Reptiles (The Reptile Database - Uetz et al., 2026), Amphibians (GABiP), Mammals (Pantheria - Jones et al., 2009), Birds (AVONET - Tobias et al., 2022) |
| Genus | The taxonomic Genus assigned to said species | Character | Reptiles (The Reptile Database - Uetz et al., 2026), Amphibians (GABiP), Mammals (Pantheria - Jones |

| | | | |
|---|---|---|---|
| | | | et al., 2009), Birds (AVONET - Tobias et al., 2022) |
| Class | The taxonomic Class assigned to said species | Character | Reptiles (The Reptile Database - Uetz et al., 2026), Amphibians (GABiP), Mammals (Pantheria - Jones et al., 2009), Birds (AVONET - Tobias et al., 2022) |
| Order | The taxonomic Order assigned to said species | Character | Reptiles (The Reptile Database - Uetz et al., 2026), Amphibians (GABiP), Mammals (Pantheria - Jones et al., 2009), Birds (AVONET - Tobias et al., 2022) |
| Family | The taxonomic Family assigned to said species | Character | Reptiles (The Reptile Database - Uetz et al., 2026), Amphibians (GABiP), Mammals (Pantheria - Jones et al., 2009), Birds (AVONET - Tobias et al., 2022) |
| Mid_Genome | The median genome size (c-value) calculated for said species | Numeric continuous | Animal Genome Database (Gregory, 2025) and National Library of Medicine (NCBI, 2025), and GABiP for amphibians |
| invaded.Africa | Record of an impact occurring due to | Binary | GIDIAS - Bacher et al., 2025 |

| | | | |
|---|---|---|---|
| | the an INNS in the region ‘Africa’ | | |
| invaded.Europe_CentralAsia | Record of an impact occurring due to the INNS in the region ‘Europe and Central Asia’ | Binary | GIDIAS - Bacher et al., 2025 |
| invaded.Asia_Pacific | Record of an impact occurring due to the INNS in the region ‘Asia and Pacific Islands’ | Binary | GIDIAS - Bacher et al., 2025 |
| invaded.Antarctica | Record of an impact occurring due to the INNS in the region ‘Antarctica’ | Binary | GIDIAS - Bacher et al., 2025 |
| invaded.Americas | Record of an impact occurring due to the INNS in the region ‘Americas’ | Binary | GIDIAS - Bacher et al., 2025 |
| Island.invaded | Record of an impact occurring due to the INNS on an island | Binary | GIDIAS - Bacher et al., 2025 |
| PA.invaded | Record of an impact occurring due to the INNS in a recognised protected area (PA) | Binary | GIDIAS - Bacher et al., 2025 |
| invaded.Freshwater | Record of an impact occurring due to the INNS in a freshwater habitat | Binary | GIDIAS - Bacher et al., 2025 |
| invaded.Marine | Record of an impact occurring due to the INNS in a marine habitat | Binary | GIDIAS - Bacher et al., 2025 |

| | | | |
|---|---|---|---|
| invaded.Terrestrial | Record of an impact occurring due to the INNS in a terrestrial habitat | Binary | GIDIAS - Bacher et al., 2025 |
| local.spatial.scale | Whether the spatial scale of the impact is ‘local’ | Binary | GIDIAS - Bacher et al., 2025 |
| global.spatial.scale | Whether the spatial scale of the impact is ‘global’ | Binary | GIDIAS - Bacher et al., 2025 |
| Nature.negative.impact | Whether the INNS has a negative impact on nature | Binary | GIDIAS - Bacher et al., 2025 |
| Nature.neutral.impact | Whether the INNS has a neutral impact on nature | Binary | GIDIAS - Bacher et al., 2025 |
| Nature.positive.impact | Whether the INNS has a positive impact on nature | Binary | GIDIAS - Bacher et al., 2025 |
| NCP.negative.impact | Whether the INNS has a negative impact on nature’s contribution to people (IPBES categories) | Binary | GIDIAS - Bacher et al., 2025 |
| NCP.neutral.impact | Whether the INNS has a neutral impact on nature’s contribution to people (IPBES categories) | Binary | GIDIAS - Bacher et al., 2025 |
| NCP.positive.impact | Whether the INNS has a positive impact on nature’s contribution to people (IPBES categories) | Binary | GIDIAS - Bacher et al., 2025 |
| Nature.direct.impact | Whether the INNS directly impact nature | Binary | GIDIAS - Bacher et al., 2025 |

| | | | |
|---|---|---|---|
| Nature.indirect.impact | Whether the INNS indirectly impact nature | Binary | GIDIAS - Bacher et al., 2025 |
| global.extinction | Whether the impacts of the INNS led to a global extinction of a native species | Binary | GIDIAS - Bacher et al., 2025 |
| competition.Nature | Whether competition was a mechanism that led the INNS to have an environmental impact | Binary | GIDIAS - Bacher et al., 2025 |
| Indirect.interactions.Nature | Whether indirect interactions were a mechanism that led the INNS to have an environmental impact | Binary | GIDIAS - Bacher et al., 2025 |
| predation.Nature | Whether predation was a mechanism that led the INNS to have an environmental impact | Binary | GIDIAS - Bacher et al., 2025 |
| herbivory.Nature | Whether herbivory was a mechanism that led the INNS to have an environmental impact | Binary | GIDIAS - Bacher et al., 2025 |
| hybrid.Nature | Whether hybridisation was a mechanism that led the INNS to have an environmental impact | Binary | GIDIAS - Bacher et al., 2025 |
| disease.Nature | Whether disease spread was a | Binary | GIDIAS - Bacher et al., 2025 |

| | | | |
|---|---|---|---|
| | mechanism that led the INNS to have an environmental impact | | |
| poisoning.Nature | Whether poisoning was a mechanism that led the INNS to have an environmental impact | Binary | GIDIAS - Bacher et al., 2025 |
| CPS.impact.Nature | Whether chemical, physical or structural impacts were mechanisms that led the INNS to have an environmental impact | Binary | GIDIAS - Bacher et al., 2025 |
| parasitism.Nature | Whether parasitism was a mechanism that led the INNS to have an environmental impact | Binary | GIDIAS - Bacher et al., 2025 |
| biofoul.Nautre | Whether biofouling was a mechanism that led the INNS to have an environmental impact | Binary | GIDIAS - Bacher et al., 2025 |
| Ecological_impact | Whether this species has a negative ecological impact | Binary | RAMA |
| InvaCost_Urban_invasion | Whether the economic impact occur in an urban area | Binary | InvaCost - Diagne et al., 2020a |
| InvaCost_Island_invaded | Whether the economic impacts occur on an island | Binary | InvaCost - Diagne et al., 2020a |
| InvaCost_PA_invaded | Whether the economic | Binary | InvaCost - Diagne et al., 2020a |

| | | | |
|---|---|---|---|
| | impacts occur in a protected area | | |
| inva_invaded_Africa | Whether the economic impacts occur in Africa | Binary | InvaCost - Diagne et al., 2020a |
| inva_invaded_AntarcticSubantarctic | Whether the economic impacts occur in Antarctic or Subantarctic | Binary | InvaCost - Diagne et al., 2020a |
| inva_invaded_Asia | Whether the economic impacts occur in Asia | Binary | InvaCost - Diagne et al., 2020a |
| inva_invaded_CentralAmerica | Whether the economic impacts occur in central America | Binary | InvaCost - Diagne et al., 2020a |
| inva_invaded_Europe | Whether the economic impacts occur in Europe | Binary | InvaCost - Diagne et al., 2020a |
| inva_invaded_NorthAmerica | Whether the economic impacts occur in North America | Binary | InvaCost - Diagne et al., 2020a |
| inva_invaded_Oceania | Whether the economic impacts occur in Oceania | Binary | InvaCost - Diagne et al., 2020a |
| inva_invaded_PacificIslands | Whether the economic impacts occur across the Pacific islands | Binary | InvaCost - Diagne et al., 2020a |
| inva_invaded_SouthAmerica | Whether the economic impacts occur in South America | Binary | InvaCost - Diagne et al., 2020a |
| inva_impacted_Agriculture | Whether the economic impacts affect the agricultural sector | Binary | InvaCost - Diagne et al., 2020a |
| inva_impacted_AuthoritiesStakeholders | Whether the economic | Binary | InvaCost - Diagne et al., 2020a |

| | | | |
|---|---|---|---|
| | impacts affect the authorities and stakeholders sector | | |
| inva_impacted_Environment | Whether the economic impacts affect the environmental sector | Binary | InvaCost - Diagne et al., 2020a |
| inva_impacted_Fishery | Whether the economic impacts affect the fishery sector | Binary | InvaCost - Diagne et al., 2020a |
| inva_impacted_Forestry | Whether the economic impacts affect the forestry sector | Binary | InvaCost - Diagne et al., 2020a |
| inva_impacted_Health | Whether the economic impacts affect the health sector | Binary | InvaCost - Diagne et al., 2020a |
| inva_impacted_PublicSocialWelfare | Whether the economic impacts affect the public social welfare sector | Binary | InvaCost - Diagne et al., 2020a |
| inva_CostType_Damage | Whether the monetary cost of the INNS due to damage caused by the species | Binary | InvaCost - Diagne et al., 2020a |
| inva_CostType_Management | Whether the monetary cost of the INNS due to the management of the species | Binary | InvaCost - Diagne et al., 2020a |
| inva_ManagmentType_Knowledge | Whether the management cost for this species fall under ‘knowledge funding’ | Binary | InvaCost - Diagne et al., 2020a |

| | | | |
|---|---|---|---|
| inva_ManagmentType_PostInvasion | Whether the management cost for this species occur post-invasion | Binary | InvaCost - Diagne et al., 2020a |
| inva_ManagmentType_PreInvasion | Whether the management cost for this species occur pre-invasion | Binary | InvaCost - Diagne et al., 2020a |
| Economic_impact | Whether this INNS have a recorded monetary cost | Binary | RAMA |
| Economic.Ecological_impact | Whether this species have an economic and ecological impact | Binary | RAMA |
| Establishment_Status | Introduced status of the taxon in the country | Binary | GRIIS - Pagad et al., 2018 |
| InvaCost_Occurrence_Onetime | Whether the economic impact of the INNS a one-time occurrence | Binary | InvaCost - Diagne et al., 2020a |
| isInvasive | Invasive status; whether this species' introduction and/or spread threatens biological diversity | Categorical | GRIIS - Pagad et al., 2018 |
| Economic_impact_binary | Whether this INNS has a recorded monetary cost | Binary | RAMA |
| Economic.Ecological_impact_binary | Whether this species has an economic and ecological impact | Binary | RAMA |
| establishmentMeans | Introduced status of the taxon in the country | Categorical | GRIIS - Pagad et al., 2018 |

| | | | |
|---|---|---|---|
| num_countries_invaded | The number of countries the INNS is established within | Numeric continuous | GRIIS - Pagad et al., 2018 |
| num_continents_invaded | The number of continents the INNS is established within | Numeric continuous | GRIIS - Pagad et al., 2018 |
| Mean_Cost | The average cost of INNS impact from all recorded invasion costs | Numeric continuous | InvaCost - Diagne et al., 2020a |
| Total_Cost | The total recorded cost of INNS impact from all recorded invasion costs | Numeric continuous | InvaCost - Diagne et al., 2020a |
| first_impact_year | The earliest recorded year of an ecological impact from this INNS | Numeric continuous | GIDIAS - Bacher et al., 2025 |
| last_impact_year | The latest recorded year of an ecological impact from this INNS | Numeric continuous | GIDIAS - Bacher et al., 2025 |
| earliest_probable_starting_year | The earliest recorded year of the economic impact | Numeric continuous | InvaCost - Diagne et al., 2020a |
| mean_probable_starting_year | The mean probable starting year of the economic impact | Numeric continuous | InvaCost - Diagne et al., 2020a |
| latest_probable_ending_year | The latest recorded year of the economic impact | Numeric continuous | InvaCost - Diagne et al., 2020a |
| InvaCost_Occurrence_P_Ongoing | The economic impact of the INNS ongoing | Binary | InvaCost - Diagne et al., 2020a |
| mean.magnitude.nature.impact | According to the EICAT impact magnitudes, where does the | Numeric count | GIDIAS - Bacher et al., 2025 |

| | species impact sit between 1 and 5? |
|---|---|

***Table S3.*** *The outputs of the chi squared post hoc test, providing the Chi-square, p-unadjusted and p-adjusted values for each pairwise comparison of genome size availability between the four tetrapod taxonomic classes from within the Reptilia, Amphibia, Mammalia and Aves.*

| *Comparison* | *Chi squared test coefficients* | | |
|---|---|---|---|
| | ChiSq | Raw p | P.adj |
| *Amphibia - Aves* | **594.36** | **< 0.001** | **< 0.001** |
| *Amphibia - Mammalia* | **637.05** | **< 0.001** | **< 0.001** |
| *Aves - Mammalia* | **105.5** | **< 0.001** | **< 0.001** |
| *Amphibia - Reptilia* | **11.56** | **< 0.001** | **< 0.001** |
| *Aves - Reptilia* | **1397.38** | **< 0.001** | **< 0.001** |
| *Mammalia - Reptilia* | **1448.02** | **< 0.001** | **< 0.001** |

***Table S4***. *The outputs of the pairwise comparison post hoc Dunn's test, providing z, p-unadjusted and p-adjusted values for each pairwise comparison of genome size between the four tetrapod taxonomic classes from within the Reptilia, Amphibia, Mammalia and Aves.*

| *Comparison* | *Dunn's Post hoc test coefficients* | | |
|---|---|---|---|
| | z | P.unadj | P.adj |
| *Amphibia - Aves* | **40.26** | **0.00** | **0.00** |
| *Amphibia - Mammalia* | **8.67** | **< 0.001** | **< 0.001** |
| *Aves - Mammalia* | **-42.43** | **0.00** | **0.00** |
| *Amphibia - Reptilia* | **12.83** | **< 0.001** | **< 0.001** |
| *Aves - Reptilia* | **-19.34** | **< 0.001** | **< 0.001** |
| *Mammalia - Reptilia* | **7.07** | **< 0.001** | **< 0.001** |

***Table S5***. *The outputs of the chi squared and chi squared post hoc test, providing the Chi-square, p-unadjusted and p-adjusted values for each pairwise comparison of establishment record occurrence between the four tetrapod taxonomic classes from within the Reptilia, Amphibia, Mammalia and Aves.*

| *Comparison* | *Chi squared test coefficients* | | |
|---|---|---|---|
| | ChiSq | Raw p | P.adj |
| *Amphibia - Aves* | **317.31** | **< 0.001** | **< 0.001** |
| *Amphibia - Mammalia* | **155.2** | **< 0.001** | **< 0.001** |
| *Aves - Mammalia* | **46.92** | **< 0.001** | **< 0.001** |
| *Amphibia - Reptilia* | **20.98** | **< 0.001** | **< 0.001** |
| *Aves - Reptilia* | **206.76** | **< 0.001** | **< 0.001** |
| *Mammalia - Reptilia* | **51.91** | **< 0.001** | **< 0.001** |

***Table S6***. *The outputs of the pairwise comparison post hoc Dunn's test, providing z, p-unadjusted and p-adjusted values for each pairwise comparison of number of countries invaded between the four tetrapod taxonomic classes from within the Reptilia, Amphibia, Mammalia and Aves.*

| *Comparison* | *Dunn's Post hoc test coefficients* | | |
|---|---|---|---|
| | z | P.unadj | P.adj |
| *Amphibia - Aves* | -1.41 | 0.16 | 0.96 |
| *Amphibia - Mammalia* | -1.81 | 0.07 | 0.42 |
| *Aves - Mammalia* | -0.89 | 0.37 | 1.00 |
| *Amphibia - Reptilia* | 0.82 | 0.41 | 1.00 |
| *Aves - Reptilia* | **3.9** | **< 0.001** | **< 0.001** |
| *Mammalia - Reptilia* | **3.89** | **< 0.001** | **< 0.001** |

***Table S7***. *The outputs of the pairwise comparison post hoc Dunn's test, providing z, p-unadjusted and p-adjusted values for each pairwise comparison of the mean impact magnitude scores between the four tetrapod taxonomic classes from within the Reptilia, Amphibia, Mammalia and Aves.*

| *Comparison* | *Dunn's Post hoc test coefficients* | | |
|---|---|---|---|
| | z | P.unadj | P.adj |
| *Amphibia - Aves* | 2.33 | 0.02 | 0.12 |
| *Amphibia - Mammalia* | -1.94 | 0.05 | 0.31 |
| *Aves - Mammalia* | **-5.64** | **<0.001** | **<0.001** |
| *Amphibia - Reptilia* | -1.09 | 0.27 | 1.00 |
| *Aves - Reptilia* | **-3.07** | **0.002** | **0.01** |
| *Mammalia - Reptilia* | 0.3 | 0.76 | 1.00 |

***Table S8.*** *The outputs of the fisher's exact post hoc test, providing the Chi-square, p-unadjusted and p-adjusted values for each pairwise comparison of impact record distribution across the four tetrapod taxonomic classes from within the Reptilia, Amphibia, Mammalia and Aves.*

| *Comparison* | *Chi squared test coefficients* | | |
|---|---|---|---|
| | ChiSq | Raw p | P.adj |
| *Amphibia - Aves* | **317.31** | **< 0.001** | **< 0.001** |
| *Amphibia - Mammalia* | **155.2** | **< 0.001** | **< 0.001** |
| *Aves - Mammalia* | **46.92** | **< 0.001** | **< 0.001** |
| *Amphibia - Reptilia* | **20.98** | **< 0.001** | **< 0.001** |
| *Aves - Reptilia* | **206.76** | **< 0.001** | **< 0.001** |
| *Mammalia - Reptilia* | **51.91** | **< 0.001** | **< 0.001** |

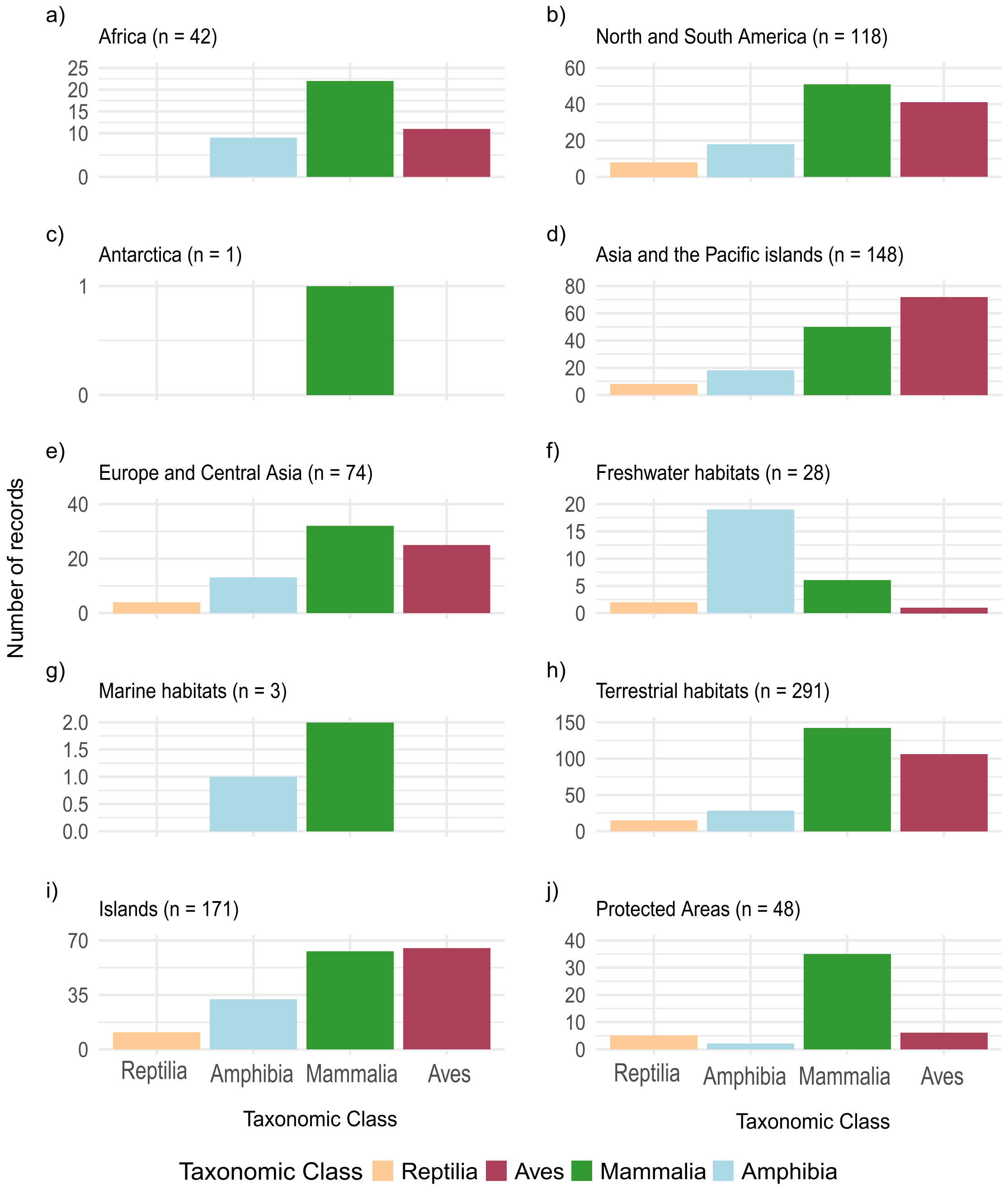

***Figure S2*** *– The distribution and number of invasive alien species (INNS) across geographic regions (a-e) and habitat types (f-j). The geographic regions include: a) Africa, b) North and South America, c) Antarctica, d) Asia and the Pacific islands, and e) Europe and Central Asia). The habitat types include: f) Freshwater, g) Marine, h) Terrestrial, i) Islands, and j) Protected Areas). INNS abundance has been displayed across the taxonomic classes – Reptilia, Amphibia, Mammalia and Aves.*

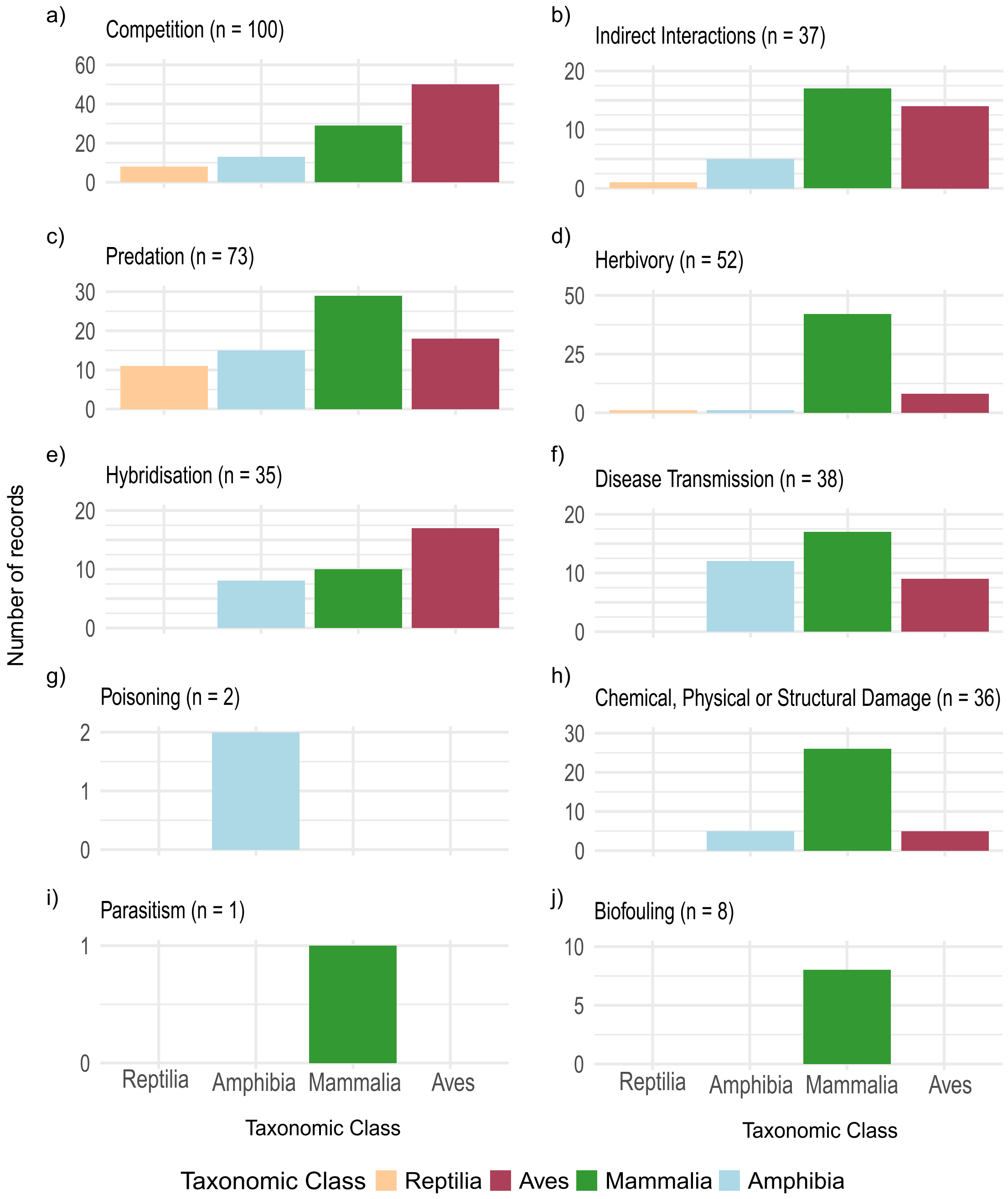

***Figure S3*** *– The frequency of the 'usage' of invasion impact mechanisms by Reptilia, Amphibia, Mammalia and Aves invasive alien species. The mechanisms include: a) Competition, b) Indirect Interactions, c) Predation, d) Herbivory, e) Hybridisation, f) Disease Transmission, g) Poisoning, h) Chemical, Physical or Structural Damage, i) Parasitism, and j) Biofouling.*
